# Astrocyte and neuronal Panx1 support long-term reference memory in mice

**DOI:** 10.1101/2023.01.16.524236

**Authors:** Price Obot, Galadu Subah, Antonia Schonwald, Jian Pan, Libor Velíšek, Jana Velíšková, Patric K. Stanton, Eliana Scemes

## Abstract

Pannexin 1 (Panx1) are ubiquitously expressed proteins that form plasma membrane channels permeable to anions and moderate sized signaling molecules (e.g., ATP, glutamate). In the nervous system, activation of Panx1 channels have been extensively shown to contribute to distinct neurological disorders (epilepsy, chronic pain, migraine, neuroAIDS, etc.) but knowledge of extent to which these channels have a physiological role remains restricted to three studies supporting their involvement in hippocampus dependent learning. Given that Panx1 channels may provide an important mechanism for activity-dependent neuron-glia interaction, we used Panx1 transgenic mice with global and cell-type specific deletions of Panx1 to interrogate their participation in working and reference memory. Using the 8-arm radial maze, we show that long-term spatial reference memory, but not spatial working memory, is deficient in Panx1-null mice and that both astrocyte and neuronal Panx1 contribute to the consolidation of long-term spatial memory. Field potential recordings in hippocampal slices of Panx1-null mice revealed an attenuation of both long-term potentiation (LTP) of synaptic strength and long-term depression (LTD) at Schaffer collateral – CA1 synapses without alterations basal synaptic transmission or pre-synaptic paired-pulse facilitation. Our results implicate both neuronal and astrocyte Panx1 channels as critical players for the development and maintenance of long-term spatial reference memory in mice.

## INTRODUCTION

Pannexins (Panxs) are a group of proteins that belong to the gap junction family but do not form gap junction channels. Of the three pannexins (Panx1, Panx2 and Panx3), Panx1 is the best characterized, with studies indicating that this protein forms plasma membrane channels that are ubiquitously expressed in vertebrate tissues (Bruzzone et al., 2003; Hanstein et al., 2013). In the central nervous system (CNS), Panx1 is present in glia and neurons (Ray et al., 2005; Vogt et al., 2005; Santiago et al., 2011; Rigato et al., 2012; Seo et al., 2020). Panx1 channels are permeable to anions and larger negatively charged molecules such as glutamate and ATP (Ma et al., 2009; Bao et al., 2004) and are activated by voltage, caspases (Sandilos et al., 2012), and following membrane receptor activation (purinergic P2 receptors, glutamate receptors, alpha-adrenergic receptors, thromboxane receptors: (Pelegrin and Surprenant 2006; Locovei et al., 2007; Locovei et al., 2006; Weilinger et al., 2012; Billaud et al., 2011; da Silva-Souza et al., 2014). Despite the strong evidence in support of the participation of Panx1 channels in neurological disorders (Santiago et al., 2011; Thompson et al., 2006; Cisneros-Mejorado et al., 2015; Lutz et al., 2013; Velasquez et al., 2016; Dossi et al., 2018; Karatas et al., 2013; Weaver et al., 2017; Burma et al., 2017), little is known about their physiological role in the CNS. Few studies have indicated that these channels have a physiological role during neurodevelopment (Wicki-Stordeur et al., 2012; Wicki-Stordeur et al., 2016) or in the regulation of synaptic plasticity involved in learning and memory (Prochnow et al., 2012; Ardiles et al., 2014; Gajardo et al., 2018).

Evidence that Panx1 contributes to learning and memory came from studies performed on transgenic mice showing that global deletion of Panx1 impaired spatial learning, novel object recognition and spatial reversal learning (Prochnow et al., 2012; Gajardo et al., 2018). These memory related behaviors, which are dependent on hippocampal synaptic plasticity, (both long-term potentiation and depression: LTP, LTD), are altered in mice lacking Panx1 (Prochnow et al., 2012; Ardiles et al., 2014; Gajardo et al., 2018). These two major forms of synaptic plasticity require, at least in part, glutamate binding to the N-methyl-D-aspartate receptor (NMDAR) (Malenka and Bear 2004). LTP is induced by strong stimuli leading to large influx of calcium through NMDAR and activation of CaMKII, leading to persistent enhancement of synaptic strength, while weaker stimuli are believed to lead to LTD by driving smaller flux of calcium through NMDAR resulting in preferential activation of protein phosphatases leading to long-term reductions in synaptic strength (Malenka 1994; Lisman 1989). NMDARs have been linked to Panx1 activation through metabotropic signaling and Src kinase recruitment under both physiological (Bialecki et al., 2020) and pathological (Weilinger et al., 2016; Weilinger et al., 2012) conditions. Recent findings in synaptic plasticity regarding metabotropic NMDAR signaling, a calcium influx independent NMDAR signaling, (Nabavi et al., 2013; Stein et al., 2015; Stein et al., 2020) and Panx1 involvement in synaptic plasticity (Ardilles et al., 2014; Gajardo et al., 2018; Prochnow et al., 2012), suggest that the NMDAR-Src-Panx1 complex may also be involved in synaptic plasticity.

Given that Panx1 in the CNS is expressed in astrocytes and neurons (Zoidl et al., 2007; Santiago et al., 2011) and that astrocytes participate in the induction of long-term synaptic plasticity through the release of gliotransmitters, including glutamate, ATP/adenosine, and D-serine (reviewed in (Perea et al., 2009; Durkee et al., 2021)), we investigated the extent to which memory deficits reported in mice with global deletion of Panx1 can be attributed to either astrocyte or neuronal Panx1. Using behavioral tests and electrophysiological recordings from Panx1 transgenic mice, we found that Panx1 does not participate in locomotor activity or anxiety-like behavior, but that it does contribute to long-term reference memory formation. Electrophysiological recordings of hippocampal slices confirmed impairments in long-term synaptic plasticity in Panx1-null mice, reduced magnitude of both LTP and LTD, compared to wild-type control mice. Because deletion of neuronal or astrocytic Panx1 induced similar impairments of long-term reference memory as seen in the global Panx1 knockout, we concluded that both astrocyte and neuronal Panx1 are necessary for the acquisition/consolidation of long-term spatial memory.

## RESULTS

### Panx1 does not influence general locomotor activity or thigmotaxis

To evaluate the overall locomotor activity and thigmotaxis (as a proxy for anxiety-like behavior) of mice with global and cell type specific deletion of Panx1, the total distance travelled, and the percent time spent in outer and inner zones of the open field arena were measured. Given that no significant sex differences in terms of thigmotaxis were detected in Panx1^f/f^ or Panx1-null mice (Supplemental Figure 1S), subsequent behavioral analysis was performed by grouping male and female mice. There was no significant difference in total distance travelled during the 10 min test between the four mouse genotypes (Figure 1A; Kruskal-Wallis, p = 0.8313). Figure1B shows the percent time that the animals spent in the outer zone; mice of all genotypes spent 92 to 96% time in the outer zone (Kruskal-Wallis, p = 0.4739). These results show that Panx1 does not affect overall locomotor activity and does not contribute to anxiety-like behavior as measured by thigmotaxis (time spent in the outer zone).

**Figure 1.**
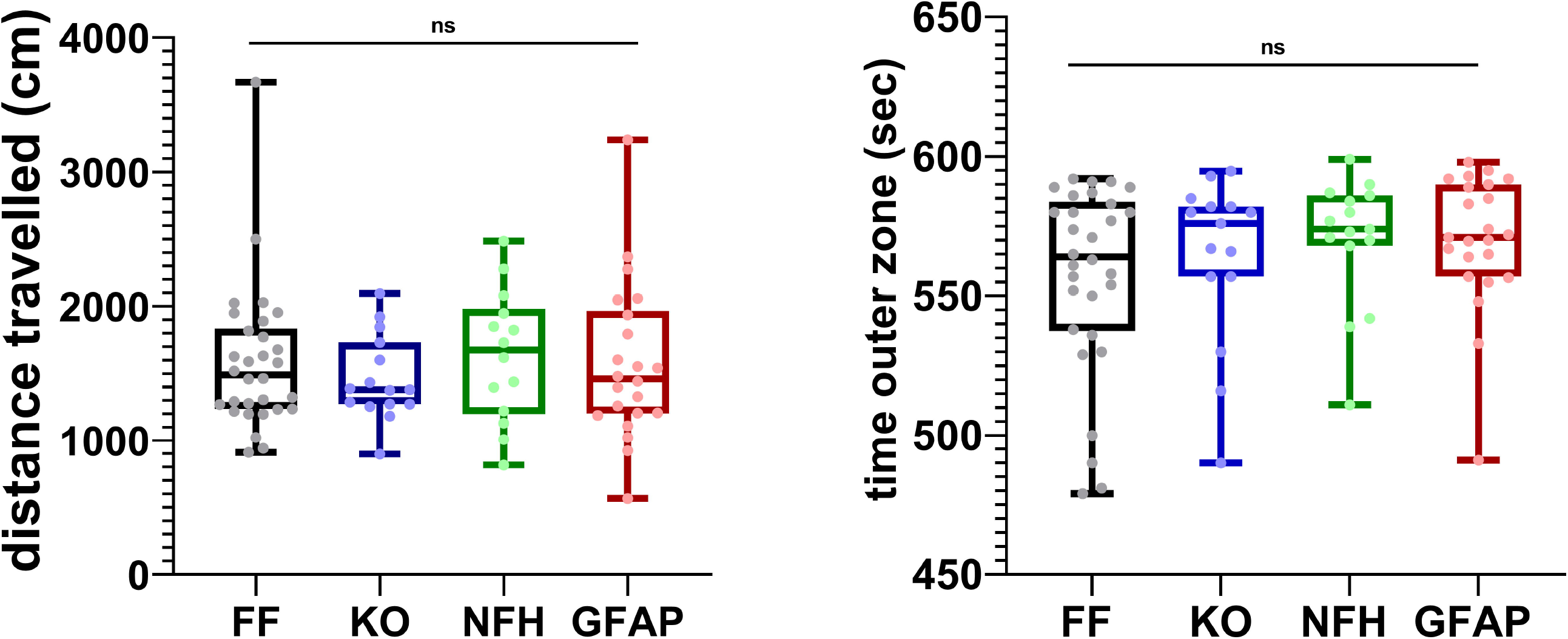
Locomotor activity and thigmotaxis are independent of Panx1. Median ± min/max values of the (**A**) total distance travelled and (**B**) time spent in the outer zone of the arena recorded from Panx1^f/f^ (FF, black bars; n = 30), Panx1-null (KO, blue bars; n = 15), NFH-Cre:Panx1^f/f^ (NFH, green bars; n =14), and GFAP-Cre:Panx1^f/f^ (GFAP, red bars; n = 23 mice) mice exposed for 10 min to the open field. ns: not significant (Kruskal-Wallis, P = 0.8313 (distance) and P = 0.4739 (time in outer zone).

### Long-Term Spatial Reference Memory is impaired in mice deficient in Panx1

The contribution of Panx1 to spatial memory was assessed using the 8-arms radial maze that allows for the simultaneous evaluation of working and reference memories. Spatial working memory is a short-term storage of task-relevant information to guide the ongoing and upcoming behavior, i.e., a temporally based representation of a spatial location that is used within a single trial of an experimental session to guide behavior. Spatial reference memory is a form of long-term memory representing the spatial, contextual, and factual aspects of a task that remains constant between trials, thus it is the representation of an association between spatial locations that remains consistent across several trials of an experimental session and is used to guide behavior.

Working and reference memory errors in the 8-arm radial maze were measured daily over 13 days during the choice phase of the training sessions (see Methods). As for thigmotaxis (Supplemental Figure 1S), no sex differences were detected in terms of working memory errors (WME) and reference memory errors (RME) in Panx1^f/f^ or Panx1-null mice (Supplemental Figure 2S). In terms of working memory, no significant differences were detected among the genotypes during the training and test sessions. (Figure 2; Supplemental Figure 3S). Figures 2A, C show that during the 13 days’ training sessions, the fraction of working memory correct (WM-C) and working memory incorrect (WM-I) errors decreased significantly over time for each mouse genotype (repeated measure ANOVA; F_time_ (12, 660) = 4.396, p < 0.0001, and F_time_ (12, 660) = 4.537, p < 0.0001, respectively), indicating that mice learned the task by reducing the number of errors. In addition, during the training sessions, no significant differences in terms of WM-C and WM-I were detected between genotypes (Figures. 2A, C; F_genotype_ (3, 55) = 0.9708, p = 0.4132, and F_genotype_ (3, 55) = 2.290, p = 0.0885, respectively). In terms of absolute number of WM errors, the number of WM-I measured for each mouse genotype was higher than the number of WM-C (Supplemental Figure S). At day 14 (test day), WM-C and WM-I did not differ significantly among the genotypes (Figures. 2B, D; Kruskal-Wallis, p = 0.2940, and p = 0.2598, respectively). Thus, these results show that independently of the genotype, mice improved their working memory performance over time and that Panx1 does not play a role, at least in this form of learning.

**Figure 2.**
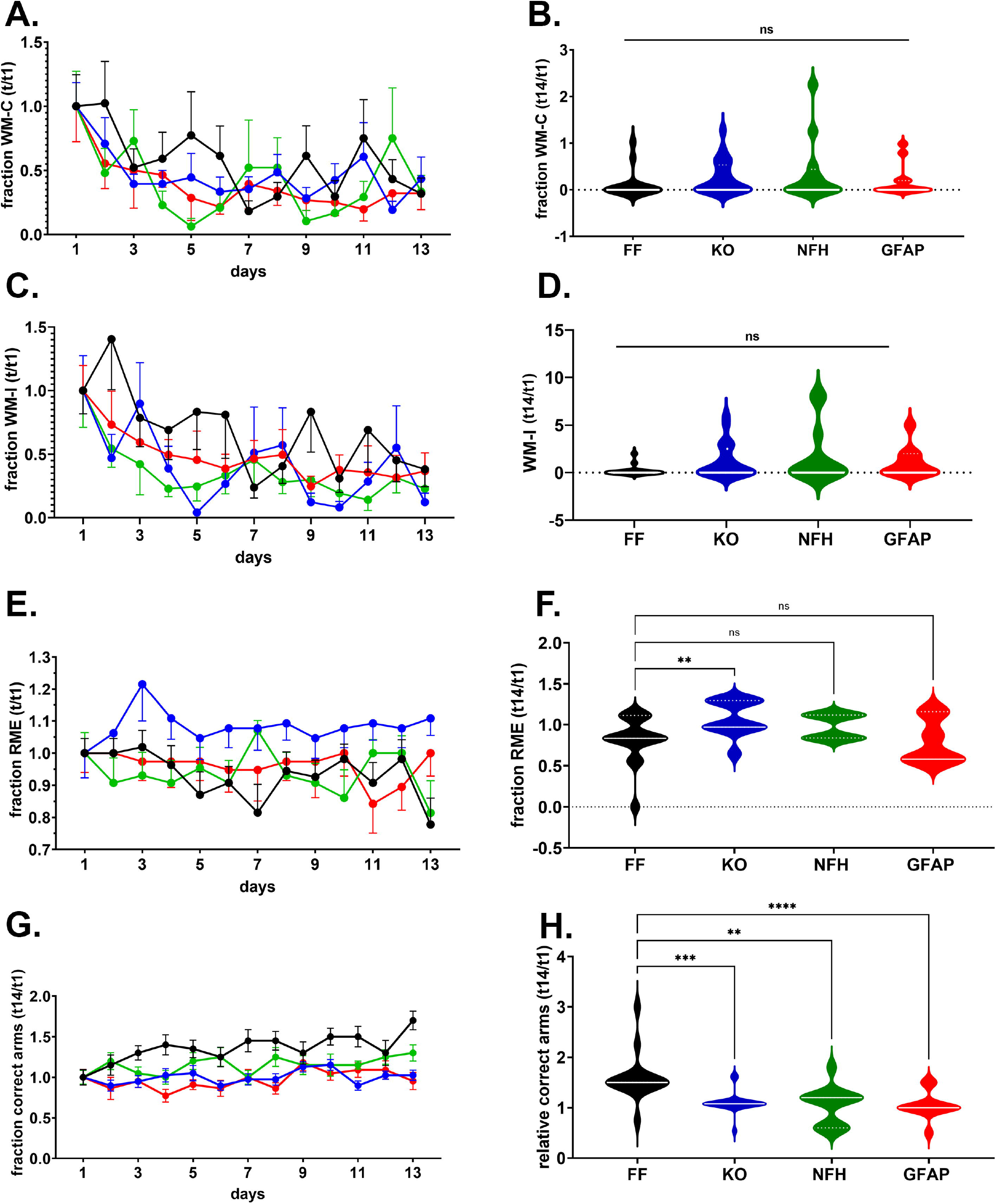
Impaired long-term reference memory in mice lacking Panx1. (**A, C, E, G**) Mean ± sem of the (**A**) fraction of working memory correct errors (WM-C), (**C**) working memory incorrect errors (WM-I), (**E**) reference memory errors (RME), and (**G**) correct choices (correct arms) obtained during the training sessions. (**B, D, F, H**) Violin plot showing the median values of WM-C (**B**), WM-I (**D**), RME (**F**), correct arms (**H**) obtained during the test phase for Panx1^f/f^ (FF, black symbols; n = 15), Panx1-null (KO, blue symbols; n = 21), NFH-Cre:Panx1^f/f^ (NFH, green symbols; n = 12), and GFAP-Cre:Panx1^f/f^ (GFAP, red symbols; n = 11) mice. ns: not significant, ** P<0.01, *** P<0.001, **** P<0.0001 (Kruskal-Wallis followed by Dunn’s multiple comparison tests).

Differently from WME, the fraction of RME recorded during the training sessions did not vary overtime within each genotype (two-ways ANOVA F_time_ (12, 660) = 0.5277, p = 0.8975), indicating that mice did not significantly improve their performance during the training phase (the number of errors did not change; see Supplemental Figure 3S). However, there was a significant difference in RME among the genotypes (Figure 2E; repeated measures ANOVA: F_genotype_ (3, 55) = 11.27, p < 0.0001), particularly between Panx1^f/f^ and Panx1-null mice (Sidak’s multi-comparison test: P < 0.0001) that was maintained during the test session, (Figure 2F; Panx1^f/f^: 0.8 folds, Panx1-null: 1.1 folds; Kruskal-Wallis followed by Dunn’s multiple comparison test: p = 0.0009).

In terms of fraction of correct (baited) arms first visited, significant differences were recorded during the 13 days of training sessions (Figure 2G; repeated measures ANOVA: F_time_ (12, 660) = 2.253, p = 0.0085) and among the genotypes (F_genotype_ (3, 55) = 56.44, p < 0.0001). At day 14 (test session), a significant deficit in long-term reference memory, in terms of correct entries into baited arms, was detected in the three genotypes that were Panx1 deficient (Figure 2H). The fold changes (mean ± sem) in the number of correct entries relative to day 1 recorded for Panx1-null (1.10 ± 0.05 fold, n = 20 mice), NFH-Cre:Panx1^f/f^ (1.05 ± 0.11 fold, n = 12 mice), and GFAP-Cre:Panx1^f/f^ (1.05 ± 0.08 fold, n = 11 mice) were significantly lower than those recorded in Panx1^f/f^ mice (1.60 ± 0.12 fold, n = 15 mice) (Figure 2H; Kruskal-Wallis followed by Dunn’s test; p < 0.01). No significant difference in terms of fraction of correct entries were detected between Panx1-null, NFH-Cre:Panx1^f/f^ and GFAP-Cre:Panx1^f/f^ mice during the test session (Dunn’s test, p > 0.55).

Thus, these results show that Panx1 contributes to long-term reference memory and that deletion of Panx1 from either astrocytes or neurons results in similar memory deficits, at least in terms of correct choices, as seen in the global Panx1-null.

### Panx1 does not contribute to basal synaptic transmission

To assess potential changes in excitatory synaptic transmission and cell excitability that could underlie the altered behavior in Panx1-deficient mice, we recorded field excitatory postsynaptic potentials (fEPSP) in response to different stimulation intensities in the apical dendritic region of *stratum radiatum* of field CA1 in hippocampal slices from young adult (2-month-old) male and female Panx1-null and Panx1^f/f^ mice. Given that no sex differences were detected when performing behavioral studies (see above), electrophysiological data derived from slices of female and male mice were pooled together. As shown in Figure 3A, no significant differences in fEPSP slopes as a function of stimulus intensity (20 – 200 μA) were detected between the two genotypes (two-way ANOVA: F_genotype_ (1, 400) = 0.0003, p = 0.98), indicating no differences in excitatory synaptic transmission.

**Figure 3.**
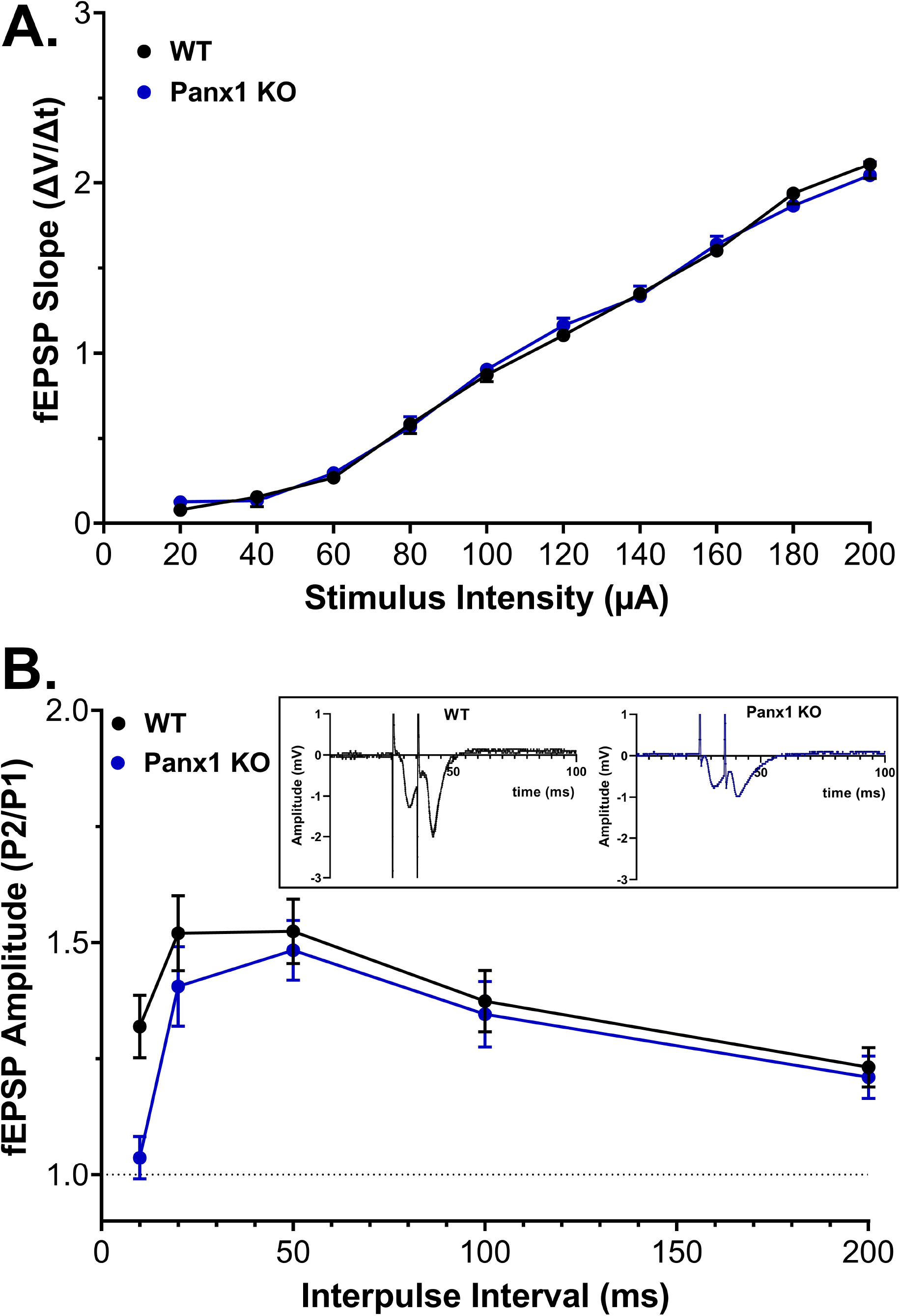
Basal synaptic transmission is unaltered in Panx1-deficient mice. (**A**) Input-output (I/O) relations showing normalized fEPSP slopes at different stimulus intensities at CA3 - CA1 synapses of hippocampal slices from control (WT) versus Panx1-null (Panx1 KO) mice. No significant difference in basal transmission was detected (Two-way ANOVA: F (9, 400) = 0.3184, P = 0.969). (**B**) Paired-pulse facilitation of the fEPSP at various inter-stimulus intervals. No significant difference was detected between WT and Panx1 KO mice (Two-way ANOVA: F (4, 180) = 0.9580, P = 0.432). **Inset in B**: representative fEPSPs recorded from Panx1^f/f^ (WT) and Panx1-null (KO) hippocampal slices in response to a 20 ms inter-stimulus interval. WT: n = 29 slices; KO: n = 13 slices.

**Figure 4.**
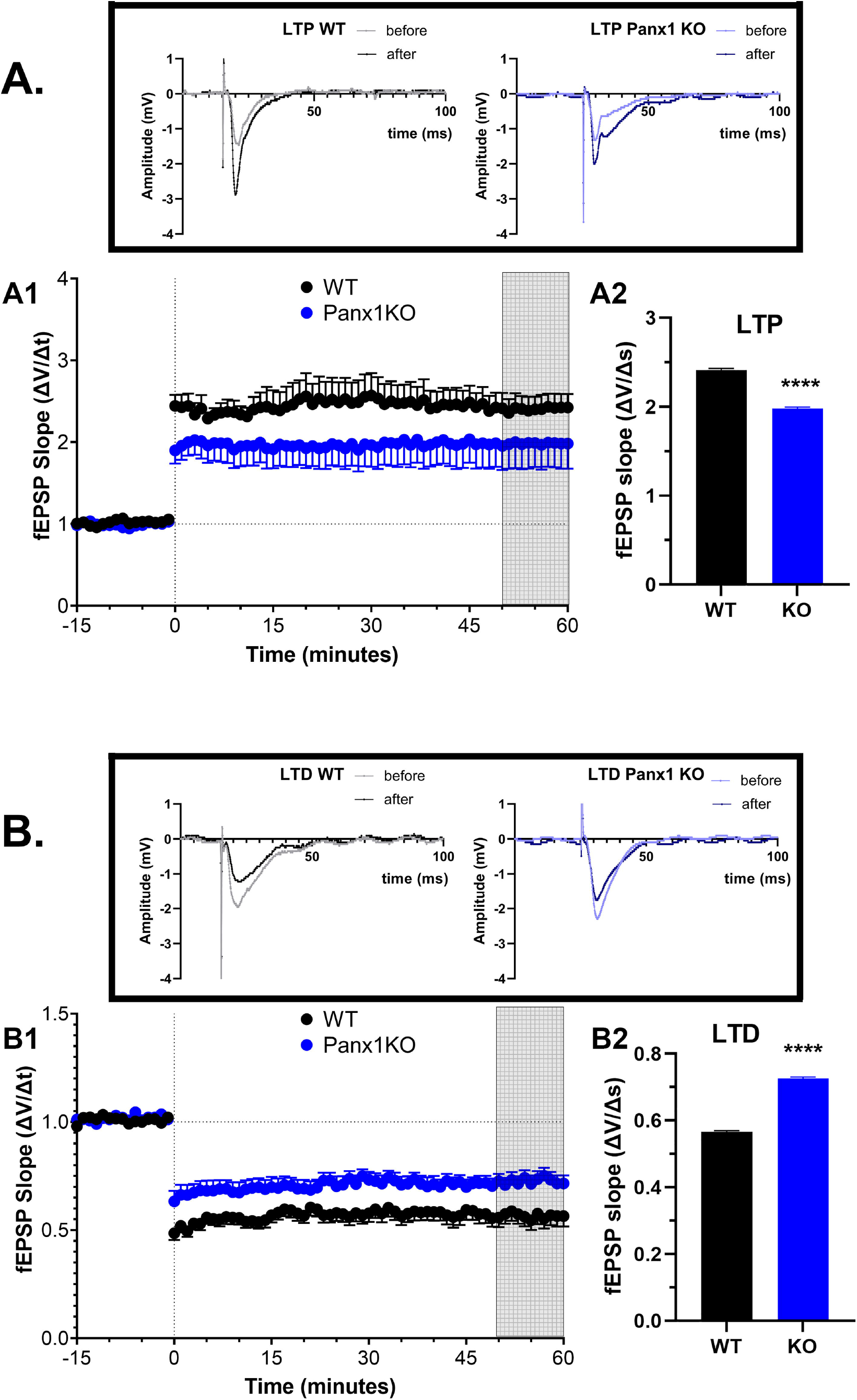
Altered hippocampal synaptic plasticity in Panx1-deficient mice. (**A-B**) Time courses (**A1, B1**) and magnitudes (**A2, B2**) of long-term potentiation (LTP) and of long-term depression (LTD). LTP was elicited by theta burst (TBS), and LTD by low frequency (LFS) stimulation, of hippocampal Schaffer collateral – CA1 synapses. Panx1-null mice (blue symbols, n = 17 slices) expressed significantly lower LTP compared to Panx1^f/f^ mice (black symbols, n = 13 slices) (Student’s t-test: t = 55.69, df = 20, p < 0.0001), and Panx1-null (blue symbols, n = 15 slices) exhibited diminished LTD compared to WT mice (black symbols, n = 12 slices) (Student’s t-test: t = 29.58, df = 20, P < 0.0001). Each data point was normalized to the averaged baseline and expressed as mean ± sem. The first set of vertical dashed lines indicate application of TBS and LFS, respectively, and the second set indicates the 50^th^ – 60^th^ minutes post-stimulation period used for LTP and LTD analyses. **Insets**: fEPSPs from representative Panx1^f/f^ (WT) and Panx1-null (KO) hippocampal slices before and after TBS for LTP **(A top)** and LFS for LTD **(B top)**.

To further characterize excitatory synaptic transmission in the hippocampal CA1 apical dendritic region in *stratum radiatum*, we measured paired-pulse facilitation (PPF: (Andersen 1960), a short-lasting form of synaptic plasticity primarily attributed to changes in presynaptic Ca^2+^ homeostasis (Zucker and Regehr 2002), with 10-200 ms inter-pulse stimulus intervals. Statistical analysis of PPF did not reveal any differences between the two genotypes (two-way ANOVA: F_genotype_ (1, 180) = 3.7687, p = 0.0538; Figure 3B).

### Panx1 contributes to hippocampal synaptic plasticity

Since deficits in long-term reference memory in spatial tasks such as the radial maze are often associated with impairments in hippocampal long-term activity-dependent synaptic plasticity (Martin et al., 2000), we sought to investigate whether the spatial learning deficits observed in Panx1 deficient mice (Figure 2) were correlated with alterations in hippocampal long-term synaptic plasticity. To that end, we examined long-term potentiation (LTP) and long-term depression (LTD) in hippocampal-dependent learning and memory in the Schaffer collateral-CA1 synapses of Panx1-null and Panx1^f/f^ mice. In slices from Panx1-null mice, the magnitude of theta burst stimulation (TBS)-induced LTP was significantly decreased compared to hippocampal slices from Panx1^f/f^ mice (Student’s t-test for unpaired data, p□<□0.0001; Figure 3A). Similar to LTP, slices from Panx1-null mice also showed impaired LTD induced by low-frequency (2 Hz/10 min) stimulation when compared to Panx1^f/f^ mice (Student’s t-test for unpaired data, p□<□0.0001; Figure 3B). There was no significant difference in 50% maximal fEPSP amplitudes recorded between the genotypes (Panx^f/f^: 72.71 ± 12.21 mV, N = 29 slices; Panx1-null: 69.26 ± 7.02 mV, N = 13 slices; Student’s t-test, p = 0.81).

These data suggest that Panx1 is a significant contributor that modulates the magnitude of LTP and LTD in young adult mice, a role that is correlated with the necessity for Panx1 for consolidation of long-term spatial reference memory in a hippocampus-dependent task.

## DISCUSSION

The present study shows that long-term spatial reference memory, but not spatial working memory, is deficient in young adult mice lacking Panx1 and that both astrocyte and neuronal Panx1 contribute to sustain long-term spatial memory. Deficits in hippocampal dependent memory in Panx1-null mice was associated with decreased long-term potentiation (LTP) and long-term depression (LTD) of synaptic transmission at Schaffer collateral – CA1 synapses, and these impairments were not related to altered neuronal excitability or presynaptic mechanisms, as indicated by input-output relations and paired pulse stimulus profiles, which were similar to those recorded from wild-type mice.

The hippocampus plays crucial roles in encoding and consolidating spatial memory (Morris et al., 1982; Squire 1992). Activity-dependent plasticity of hippocampal glutamatergic synapses, particularly NMDAR-dependent LTP and LTD, has been proposed as necessary cellular substrates for fulfilling these cognitive functions (Martin and Shapiro 2000; Bliss and Collingridge 1993; Ge et al., 2010).

The physiological contribution of Panx1 channels to synaptic plasticity, particularly to LTP and LTD, has been previously reported in older mice with global deletion of Panx1 (Prochnow et al., 2012; Ardiles et al., 2014; Gajardo et al., 2018). These studies showed that in adult (6-12 months old) mice, global deletion of Panx1 resulted in increased LTP and reduced LTD at Schaffer collateral – CA1 synapses, compared to age-matched wild-type mice (Ardiles et al., 2014; Gajardo et al., 2018). However, peripubertal Panx1-null mice (1 month old) did not display alterations in LTP and LTD (Ardilles et al., 2014). In contrast to these previous reports, our results show that both forms of long-term, activity-dependent synaptic plasticity at CA3 - CA1 synapses in hippocampus are attenuated in 2 months old Panx1-null mice. Whether these differences are age-related and/or due to the distinct genomic fabrics between the different Panx1 transgenic lines used (current study: Panx1^tm1a(KOMP)Wtsi^ -UCDavis KOMP repository; Prochnow et al., 2012: (Panx1^Vsh^ - (Romanov et al., 2012), and Ardiles et al., 2014: Panx1^-/-^ – (Anselmi et al., 2008; Bargiotas et al., 2012)) needs further investigation. Given that aberrant LTP and LTD have been associated with age-related cognitive decline (Bach et al., 1999; Barnes 1979; Burke and Barnes 2006) and that the expression of LTP-specific genes and of Panx1 declines during aging (Ardiles et al., 2014, Ryan et al., 2015, Vogt et al., 2005), it is possible that the degree by which Panx1 contributes to age-dependent synaptic plasticity will also depend on changes in LTP-specific proteins, particularly with those that Panx1 have been shown to interact/modulate, such as NMDA receptors (Bialecki et al., 2020, Weilinger et al., 2012).

Alternatively, discrepancies between our results and others may be related to differences in phenotype caused by mouse genetic background (Gerlai 1996; Simpson et al., 1997) and/or effects of passenger mutations in Panx1 congenic mice (Vanden Berghe et al., 2015). An example from a previous study found that two Panx1-null mouse lines (Panx1^Vsh^ and Panx1^Vmd^, derived respectively from 129 and C57Bl/6 embryonic stem cells) expressed the presence or absence of a phenotype that coincided with the presence or absence of one passenger mutation (Vanden Berghe et al., 2015). Thus, it is possible that the potentiation of LTP and absence of LTD found in the two 129-derived Panx1 mouse lines (Panx1^-/-^ and Panx1^Vsh^: (Ardiles et al., 2014; Prochnow et al., 2012) and the attenuation of LTP and LTD we observed for the Panx1^KOMP^ (C57BL/6-derived ECS)) could be due to the passenger mutation. Even considering this possibility, the impact of altered synaptic plasticity on the phenotype of the different mouse lines was similar, as both blockade and saturation of LTP has been found to lead to impaired spatial learning (Morris et al., 1986; Moser et al., 1998).

Our results associating reductions in both LTP and LTD with deficits in hippocampal-dependent learning are generally supported by literature implicating these forms of synaptic plasticity in spatial learning. Our findings that long-term reference, but not working, memory is impaired in Panx1 deficient mice is particularly interesting. The CA1 region of the hippocampus has been implicated in virtually all forms of hippocampus-dependent learning. However, evidence concerning its role, and that of the hippocampus at large in working memory is more limited. Highly selective genetic lesions (Brun et al., 2002) or optogenetic silencing (Nakashiba et al., 2008) of glutamatergic medial entorhinal cortex layer 3 inputs to CA1 impaired trace fear conditioning and spatial working memory, but did not perturb spatial reference memory, contextual fear conditioning, or place-cell firing. These studies allow for the proposal of dichotomous transmission pathways for working and reference memory information to CA1 pyramidal cells. That is, hippocampal-dependent reference memory may be maintained by the CA3 Schaffer collateral synaptic inputs to CA1 pyramidal neurons, whereas entorhinal cortical inputs onto apical dendrites of CA1 pyramidal neurons may play a primary role in transmitting information concerning working memory, as suggested by our results.

In light of the memory consolidation theory (Alvarez and Squire 1994), we propose that Panx1-dependent synaptic plasticity at CA1 pyramidal neurons facilitates the ability of these neurons, the major output of the hippocampus, to present or activate memories in cortical synapses for long-term storage for later retrieval. The finding that long-term reference, but not working, memory is impaired in Panx1 deficient mice is particularly interesting, suggesting that activation of Panx1 is selectively involved in the long-term plasticity in multiple brain regions that is necessary for reference memory storage.

Activity-dependent plasticity has long been emphasized as neuronal and largely synaptic alterations. However, astrocytes have been highlighted as playing important roles in the modulation of synaptic plasticity, neural circuits, and networks (Wang et al., 2021; Kastanenka et al., 2020). Besides their metabolic functions, as part of the “tripartite synapse” (Araque et al., 1999), astrocytes increase their cytoplasmic calcium levels in response to numerous synaptic signals, which induces the release of neuroactive substances (“gliotransmitters”), which in turn modulate synaptic activity and plasticity. The involvement of astrocytes in modulation of activity-dependent synaptic plasticity appears to be important for learning and memory, and thus is of behavioral relevance. For instance, astrocyte-specific genetic deletion of CB1 receptors impaired LTP and recognition memory, both of which were completely rescued by elevation of extracellular concentration of the NMDAR co-agonist D-serine (Robin et al., 2018). Moreover, a recent study showed that astrocyte p38α MAPK signaling is required for induction of LTD at CA3-CA1 synapses that contributes to contextual fear conditioning (Navarrete et al., 2019). Although more investigation will be needed to determine the mechanism(s) by which astrocyte Panx1 modulate synaptic plasticity, our data are the first to indicate that these cells have a significant role in long-term spatial reference memory. Release of ATP and D-serine through astrocytic Panx1 channels (Iglesias et al., 2009; Pan et al., 2015) represents an important mechanism for activity-dependent neuron-glia interaction. ATP acting on P2X receptors can affect the efficacy of all elements of the tripartite synapse via interaction with postsynaptic NMDA and GABA receptors, by triggering trafficking of AMPA receptors and modulating the release of neuro- and gliotransmitters (reviewed in (Pankratov et al., 2009)). D-serine stored and released from astrocytes has been shown to act on neuronal NMDARs and thus modulates NMDAR-dependent synaptic plasticity (Mothet et al., 2000; Mothet et al., 2005; Yang et al., 2003; Panatier et al., 2006; Henneberger et al., 2010).

The presence of a causal relationship between Panx1 deletion, LTP/LTD, and learning, or a correlation due to shared compensatory mechanisms, remains to be determined. Our findings that a Panx1 deletion induces impairments in the induction of both LTP and LTD in the hippocampus has significant implications for Panx1 modulation of learning and memory.

## MATERIALS and METHODS

### Animals

Panx1^f/f^ mice were generated in our facility from crosses between Panx1^tm1a(KOMP)Wtsi^ (Panx1-null; obtained from KOMP; RRID:IMSR_KOMP:CSD66379-1a-Wtsi) and a flippase deleter mouse (B6.ACTFLPe/J; RRID:MGI:5014383) in the C57BL/6 background. For astrocyte and neuronal deletion of Panx1, mGFAP-Cre (B6.Cg-Tg(Gfap-cre)73.12_Mvs/J; RRID:MGI:4829613) in the C57BL/6 background and mNFH-Cre (Tg(Nefh-cre)12_Kul/J; RRID:MGI:3043822) mice in a mixed (129/Sv * 129S7/SvEvBrd * C57BL/6 * FVB) background purchased from Jackson laboratory were crossed with Panx1^f/f^ to generate mGFAP-Cre:Panx1^f/f^ and mNFH-Cre:Panx1^f/f^ mice and maintained in our animal facility. These transgenic mice, previously described and characterized (Hanstein et al., 2013, 2016; Scemes et al., 2019), were housed in individually ventilated cages containing 3-4 males or females. Mice were located in a single room and maintained under specific pathogen-free conditions in the Animal Resource Facilities of New York Medical College and all experiments were pre-approved by the Animal Care and Use Committee.

### Behavioral Studies

For behavioral experiments 2 months old male and female mice were maintained in a light/dark cycle (light on: 7:00 AM, light off: 7:00 PM) and experiments performed during the light cycle at a time particular to each test as described below. Animals had free access to water and chow, except for two days prior to the radial maze testing when mice received a restricted diet (see below). As two-ways ANOVA showed that neither the sex nor interactions of sex with the experimental groups had any statistically significant effect (Supplemental Figures 1S and 2S), data from male and female mice were combined for behavioral analysis.

### Open Field

To determine general locomotor activity levels and thigmotaxis (tendency to remain close to the walls, and a proxy of anxiety-like behavior), we used the open field and behavioral testing performed during 9:00 AM - 12:00 PM. Animals were placed for 10 min in an arena (43.5 cm x 43.5 cm) with infra-red sensors attached to a computer. Mouse activity (total distance travelled and time spent in inner and outer zones) was recorded using Activities® software. The number of fecal boluses were determined after each mouse completed the open field test.

### Eight Arm Radial Maze

To investigate spatial orientation and memory performance in mice, we used an automated eight arm radial maze (Ugo Basile) with central circular area (16 cm) and eight arms (36 cm x 5 cm x 15 cm) that could be accessed by automated sliding doors. The maze was operated via ANY-maze video tracking software interface. The radial maze was placed on a platform (90 cm across and 38 cm on each side) which was placed directly on the floor of the testing room. External visual cues included marks on the walls near the maze, and furniture. All cues remained in the same position for all trials. Behavioral testing occurred during 1:00 – 5:00 PM. Two days before the start of the experiment, mice were put on a restricted diet and their weight maintained at 80-85% of original weight throughout the experiment. The behavioral test consisted of three sessions: (1) day one habituation session followed by (2) 13 daily training session, followed by (3) a single test session 24 hrs later. During habituation, mice were placed in the middle of the maze with access to all eight baited arms for 15 min or until the mice ate all the baits (chocolate cheerios). The baits were placed at the far end of each arm. For the training and test sessions, different sets of 4 baited arms were designated for each mouse and remained constant for any single mouse over the two weeks of training. Daily training sessions consisted of two phases, a forced and a choice phase, and lasted for 13 consecutive days. The forced phase of the training session consisted of placing the mouse on the central area for 5 sec before the 4 doors of the baited arms were simultaneously opened. After the mouse visited a baited arm, the door was closed; the forced training ended when the mouse ate all 4 baits which occurred within the 15 min forced training. The choice phase, performed 40 sec after the forced phase, consisted of placing the mouse on the central area and allowing it to enter any of the 8 (4 baited and 4 non-baited) arms. The choice phase lasted for 10 min. Twenty-four hours after the last training day, a single test session was performed. The test session was similar to the training session, but without the forced phase. The number of working and reference memory errors (see below) and the number of correct choices (number of first entries into baited arms) were recorded during the training and test sessions. Reference memory errors (RME) consisted in the first entry into a non-baited arm, while working memory errors (WME) consisted of repeated entries into previously visited arms. WME was further divided into correct (WM-C: re-entries into baited arms that have been previously visited) and incorrect (WM-I: re-entries into un-baited arms) errors, as described by others (Jarrard et al., 1984; Pirchl et al., 2010; Yan et al., 2004). For analysis, the number of WM-C, WM-I, RME and of correct choices obtained during the training and test sessions obtained for each mouse genotype were normalized to their respective mean value obtained at the first training day (day 1).

### Hippocampal slice electrophysiology

To determine the extent to which synaptic neurotransmission is modified by the genetic deletion of Panx1, electrophysiological field potential recordings from Schaffer collateral-CA1 synapses in hippocampal slices were performed, as previously described (Sullivan et al., 2018). Two-month-old Panx1^f/f^ and Panx1-null male and female mice were deeply anesthetized with isoflurane, and their brains were quickly removed and submerged in ice-cold (2–4 °C) sucrose-containing artificial cerebrospinal fluid (ACSF) (in mM): NaCl 87, NaHCO_3_ 25, NaH_2_PO_4_ 1.25, KCl 2.5, CaCl_2_ 0.5, MgCl_2_ 7, D-glucose 25 and sucrose 75), saturated with 95% O_2_/5% CO_2_. Coronal slices (350 μm) were obtained using a vibratome (Leica VT1000 S), then placed in a holding chamber containing the same sucrose-containing ACSF for at least 30 minutes before being transferred to another holding chamber containing recording ACSF (in mM: NaCl 126, NaHCO_3_ 26, NaH_2_PO_4_ 1.25, KCl 3, CaCl_2_ 2.5, MgCl_2_ 2 and D-glucose 10) for another 30 minutes. Once transferred to an interphase-type recording chamber, slices were continuously perfused with recording ACSF maintained at 32 °C. A borosilicate-glass recording electrode filled with recording ACSF (1–2 MΩ ; Model P-97, Sutter Instruments, Novato, CA, USA) was placed in the *stratum radiatum* area to record field excitatory postsynaptic potentials (fEPSPs) from distal dendritic arbors of CA1 pyramidal neurons. Evoked fEPSPs were generated by stimulating the Schaffer collaterals with pulses delivered using a bipolar stainless-steel stimulating electrode placed approximately 200-300 μm from the recording electrode, in the Schaffer collaterals/commissural fibers of *stratum radiatum*. Electrical stimulation was delivered by an ISO-Flex isolator controlled by a Master 8 pulse generator (AMPI, Jerusalem, Israel), and triggered and recorded with a Multiclamp 700B amplifier (Molecular Devices, San Jose, CA). Prior to running experimental protocols, fEPSP slopes were recorded for a 15-minute baseline. Slices exhibiting drifts larger than ±10% during baseline recording were considered not suitable and excluded from experimentation.

### Basal synaptic transmission

To evaluate for eventual changes in basic synaptic transmission caused by deletion of Panx1, we assessed neuronal excitability and synaptic transmission by measuring the input-output (I/O) relationship and pair-pulse facilitation (PPF), respectively. I/O relationship was determined from field excitatory post-synaptic potentials (fEPSP) generated at the CA1 *stratum radiatum* evoked by gradually increasing the stimulus intensity (20 μA to 200 μA; 100 μs duration; 20 s intervals) applied at the Schaffer collaterals; the input was considered as being the peak amplitude of the fiber volley and the output was the initial slope of fEPSPs. PPF was measured over 5 inter-pulse intervals of 10, 20, 50, 100, and 200 ms with intensity (50 – 100 μA) adjusted to evoke ∼50% of the maximal fEPSP. The ratio of the amplitude of the second to the first fEPSP, averaged over five sweeps per interval, was used to characterize PPF profiles.

### Long-term potentiation (LTP) and long-term depression (LTD)

A 15-minute baseline was recorded prior to experimental induction of LTP and or LTD, with the stimulus intensity of the current pulses (50 μA to 100 μA; 100 μs duration; 20 s intervals) was adjusted to evoke ∼50% of the maximal fEPSP. The high-frequency theta burst stimulus (TBS) paradigm for induction of LTP consisted of 3 theta burst trains separated by 20 seconds. Each train consisted of 10 bursts, with 5 pulses per burst, at a burst frequency of 100 Hz and inter-burst interval of 200 ms. LTD was induced using low frequency stimulation (LFS: 2 Hz/10 min). The slopes of fEPSPs were measured by linear interpolation from 20 to 80% of maximum negative deflection (SciWorks Software, DataWave Technologies, Parsippany, NJ). The magnitude of LTP and LTD was calculated as the average of the fEPSP responses recorded 50–60 min after conditioning stimulation compared to the pre-conditioning 15-minute baseline.

### Statistical Analyses

Data were analyzed and graphed using GraphPad Prism (9.4.1). The Shapiro-Wilk test was used to assess whether non-categorial data came from a normally distributed population prior to further analyses. Non-Gaussian distributed data were analyzed using the non-parametric Kruskal-Wallis’s test followed by Dunn’s multi-comparison test, and the normally distributed data analyzed using parametric test (ANOVA) when homoscedasticity was verified by visual inspection of plots of predicted Y values *versus* the absolute value of residuals. In the latter case, one-way, two-ways ANOVA with/without repeated measures followed by Sidak’s multiple comparison test were used for data analyses when appropriate. For analysis of LTP and LTD data, unpaired Student t-test was used.

## ACKOWLEDGEMENT

This work was partially supported by NIH (RO1-NS09786) to ES, JV, LV, PO.

