## Supplemental Figure S1. for "Astrocyte and neuronal Panx1 support long-term reference memory in mice"

**Supplemental Figure S1**. (**A**) Time spent in the inner zone and (**B**) number of fecal bolus recorded from male and female Panx1$^{f/f}$ (FF) and Panx1-null (KO) mice during 10 min in the open field arena. No sex and genotype differences were detected by two-way ANOVA. In parentheses are the number of mice.

**A.**

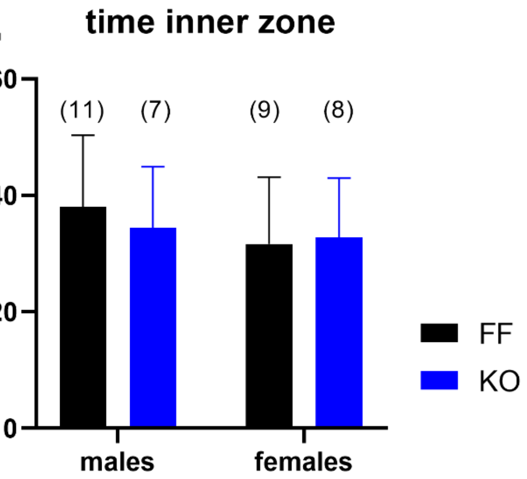


| | | |
|---|---|---|
| sex | F (1, 31) = 0.1190 | P=0.7325 |
| genotype | F (1, 31) = 0.01205 | P=0.9133 |

**B.**

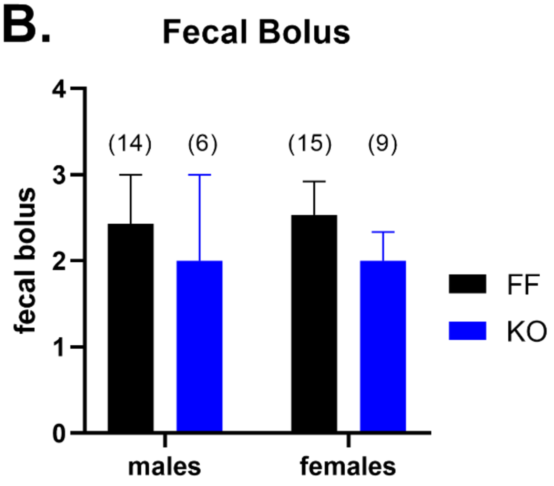


| | | |
|---|---|---|
| sex | F (1, 40) = 0.008173 | P=0.9284 |
| genotype | F (1, 40) = 0.6890 | P=0.4114 |
