## Supplemental Figure S2. for "Astrocyte and neuronal Panx1 support long-term reference memory in mice"

**Supplemental Figure S2**. Fraction of (**A**) working memory correct errors (WM-C), (**B**) working memory incorrect errors (WM-I), (**C**) reference memory errors (RME), and of (**D**) correct arms recorded from male and female Panx1$^{f/f}$ (FF) and Panx1-nul (KO) mice during the test day of the 8-arms radial maze. No significant sex differences were detected by two-way ANOVA, while significant differences in RME and correct arms were recorded between genotypes. The number of Panx1$^{f/f}$ mice used were 7 males and 8 females and of Panx1-null were 11 males and 10 females.

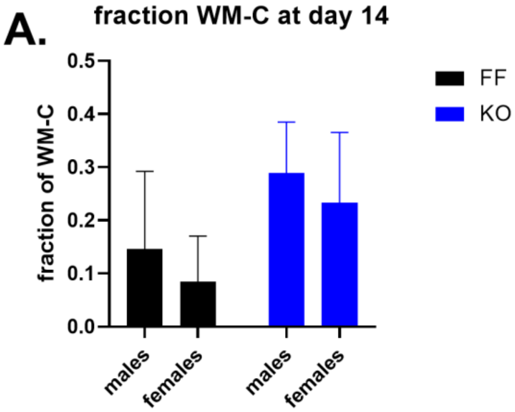


| | | |
|---|---|---|
| sex | F (1, 32) = 0.2447 | P=0.6242 |
| genotype | F (1, 32) = 1.522 | P=0.2263 |

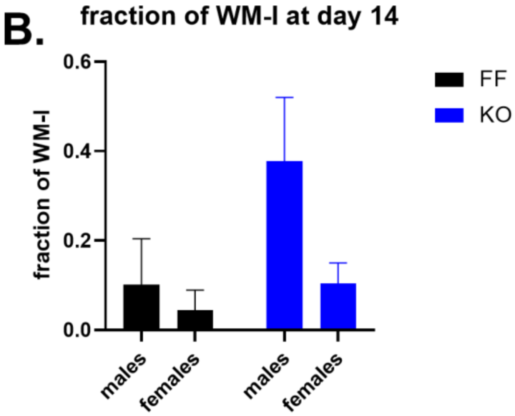


| | | |
|---|---|---|
| sex | F (1, 32) = 2.576 | P=0.1184 |
| genotype | F (1, 32) = 2.635 | P=0.1143 |

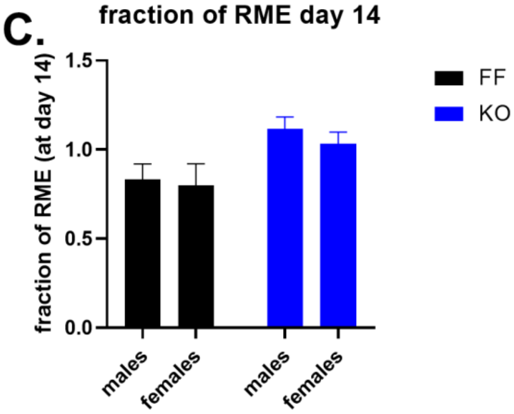


| | | |
|---|---|---|
| sex | F (1, 32) = 0.4731 | P=0.4965 |
| genotype | F (1, 32) = 9.284 | P=0.0046 |

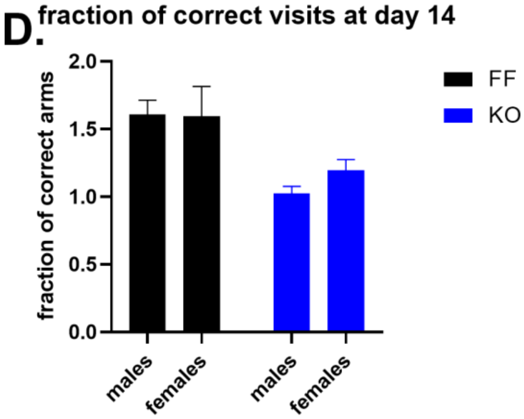


| | | |
|---|---|---|
| sex | F (1, 31) = 0.4034 | P=0.5300 |
| genotype | F (1, 31) = 15.98 | P=0.0004 |
