## Supplementary figures and images for "Astrocyte and neuronal Panx1 support long-term reference memory in mice"

### Supplemental Figure S3.

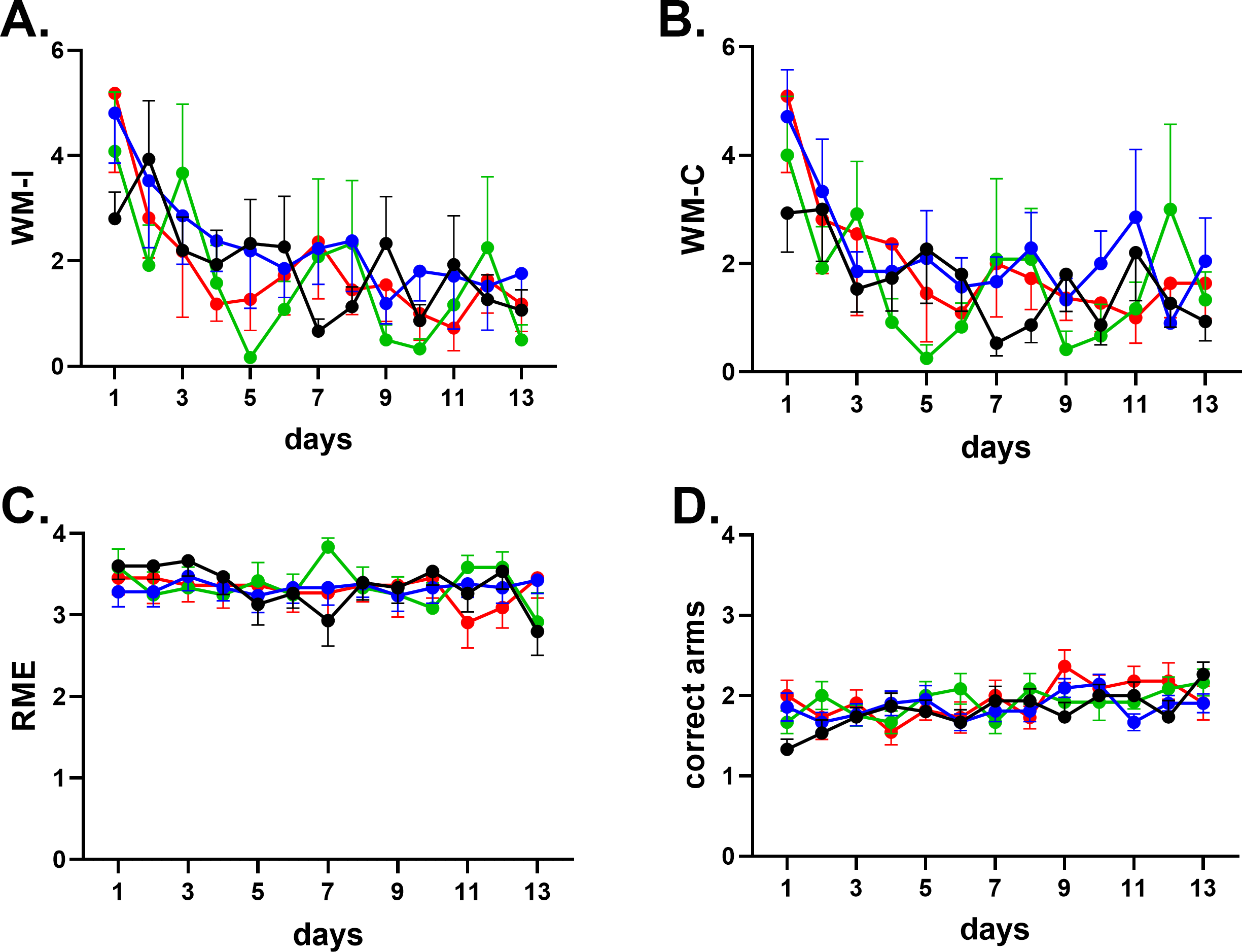

### Supplemental Figure S4.

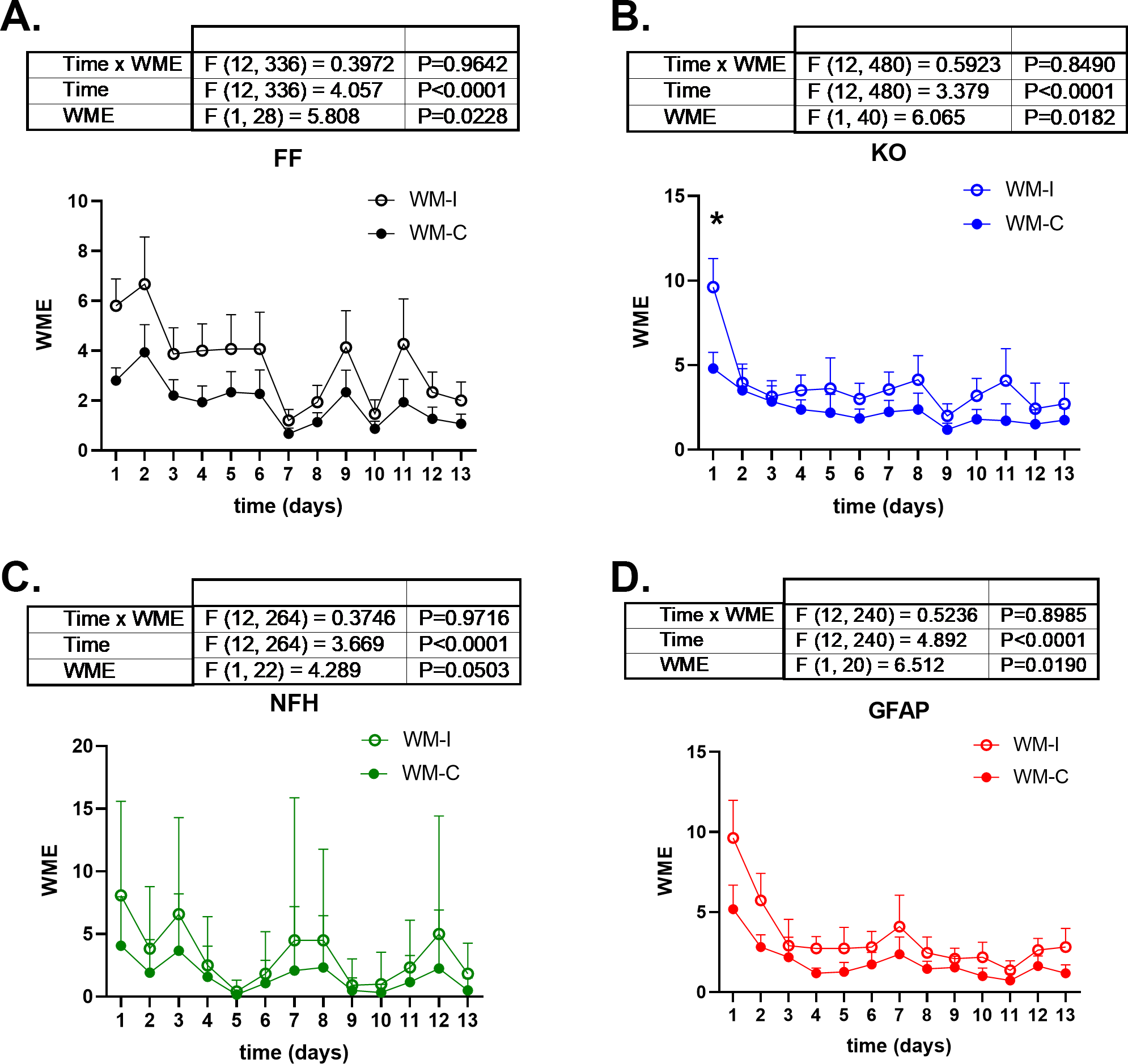
